# Rapid Sensorimotor Reinforcement in the Olfactory Striatum

**DOI:** 10.1101/730697

**Authors:** Daniel J. Millman, Venkatesh N. Murthy

**Author notes:** Allen Institute for Brain Science, Seattle, WA.

## Abstract

Rodents can successfully learn multiple, novel stimulus-response associations after only a few repetitions when the contingencies predict reward. The circuits modified during such reinforcement learning to support decision making are not known, but the olfactory tubercle (OT) and posterior piriform cortex (pPC) are candidates for decoding reward category from olfactory sensory input and relaying this information to cognitive and motor areas. Here, we show that an explicit representation for reward category emerges in the OT within minutes of learning a novel odor-reward association, whereas the pPC lacks an explicit representation even after weeks of overtraining. The explicit reward category representation in OT is visible in the first sniff (50-100ms) of an odor on each trial, and precedes the motor action. Together, these results suggest that coding of stimulus information required for reward prediction does not occur within olfactory cortex, but rather in circuits involving the olfactory striatum.

## Introduction

Reinforcement learning can quickly improve decision making by discovering and emphasizing features of the environment (e.g. sensory inputs) and actions of the animal (e.g. motor outputs) that correlate with positive or negative outcomes (Dayan and Niv, 2008; Sutton and Barto, 1998). Rodents can successfully learn to associate odor cues with motor responses following only a few repetitions of trial - and-error per odor when the correct response is reinforced by reward (Slotnick, 2001; Slotnick et al., 2000). The resulting decision process can be highly odor-specific: mice can accurately determine the presence or absence of a single rewarded odor hidden within previously unexperienced mixtures of more than a dozen distractors with similar molecular features as the target rewarded odor (Rokni et al., 2014). Converging evidence suggests that midbrain (VTA) dopaminergic neurons convey the reinforcement signal needed for learning (Cohen et al., 2012; Schultz et al., 1997), however the neural circuits that are modified through reinforcement are not known.

In the olfactory bulb, sensory processing begins with a dedicated channel (i.e. glomerulus) for each olfactory receptor. An optimal linear readout of these channels is sufficient to determine the presence of any individual odor among at least a moderately-sized (16 odor) panel of structurally similar odors and, thus, decide the appropriate motor action given a complex sensory scene (Mathis et al., 2016). The success of a linear readout implies that any monosynaptic recipient of OB output could learn to make accurate sensorimotor decisions through biologically-plausible mechanisms such as reward-gated Hebbian plasticity (Hung et al., 2005; Loewenstein and Seung, 2006; Pfeiffer et al., 2010; Reynolds and Wickens, 2002).

Although several brain regions receive input from OB, many of these regions are arranged in a feed-forward hierarchy (with abundant feedback connectivity), ultimately converging on the posterior piriform cortex (pPC) and the olfactory tubercle (OT) (Giessel and Datta, 2014; Ikemoto, 2007; Wesson and Wilson, 2011; Yamaguchi, 2017; Zhang et al., 2017a). pPC is an association cortex, with the axons of individual neurons branching and projecting to multiple cognitive brain regions, including orbitofrontal cortex, amygdala, medial temporal lobe (Diodato et al., 2016; Johnson et al., 2000). Although the OT is located in the ventral striatum (Heimer et al., 1982; Ikemoto, 2007), its anatomy more closely resembles the dorsal striatum. However, OT is specialized for olfaction with a well-defined cell body layer that receives input directly from OB in addition to input from multiple olfactory cortical areas (Wesson and Wilson, 2011; Zhang et al., 2017a). In turn, OT is by far the largest source of olfactory input to VTA (Watabe-Uchida et al., 2012), suggesting a prominent role in reward-related behaviors.

Physiological evidence suggests that an important transformation in odor coding occurs in pPC and OT during sensorimotor decision making. Activity of single-neurons that correlate with choice and reward has been observed in both pPC and OT (Calu et al., 2007; Gadziola and Wesson, 2016; Gadziola et al., 2015). The anterior piriform cortex, which immediately precedes pPC and OT in the anatomical hierarchy of olfactory regions, does not have choice activity in the first sniff of an odor stimulus (Miura et al., 2012), when decision making occurs (Abraham et al., 2004; Rinberg et al., 2006; Uchida and Mainen, 2003), although subsequent activity can reflect the decision (Gire et al., 2013). Single-neuron reward-selectivity for single odors emerges prior to the motor response at the level of OT neurons (Gadziola et al., 2015), however it is not known whether this is the case for pPC as well. If reward-selectivity is computed in pPC during decision making, OT could inherit this selectivity. Alternatively, heavy interconnection with the reward system provides the OT with the necessary reinforcement signals and plasticity-inducing dopamine for learning arbitrary odor-reward associations (Gadziola et al., 2015; Wieland et al., 2015). Indeed, responses of neurons to visual stimuli in dorsal striatum strongly reflect a newly learned reward association with a few trials of successful learning (Pasupathy and Miller, 2005; Schultz et al., 2003), so the olfactory striatum could function similarly to learn odor-outcome associations.

We developed a novel odor-reward categorization task in order to investigate the contributions of pPC and OT to odor-driven sensorimotor learning and decision making. To this end, we recorded the activity of single neurons in pPC and OT as mice learned, through trial-and-error, the reward valence assigned to a panel of previously unexperienced odor stimuli. This approach was designed to study and dissect the interaction of odor coding with reward coding in these brain regions. Along with neural recordings, we simultaneously measured sniffing in order to focus on the earliest components of sensory processing and decision making during the first sniff of odor cues. Behavioral responses indicated that mice reliably learned odor-reward contingencies within a single session, permitting the evolution of neural activity to be monitored throughout the learning process. We observed that reward selectivity was easily found in OT, multiplexed with odor selectivity, whereas explicit reward selectivity was largely absent in pPC. Together, these results support a striatal, rather than cortical, model of rodent olfactory sensorimotor learning.

## Results

### Mice quickly learn to categorize rewarded and unrewarded odors

We trained head-restrained mice to decide, based on the identity of an odor stimulus, whether to respond (Go) or not respond (No-Go) (Figure 1A; see Methods). A panel of eight odors (four rewarded and four unrewarded) was presented on randomly interleaved trials during each experimental session. This panel size enabled us to characterize the sparseness of odor tuning while yielding 20-30 repetitions per odor to accurately measure behavioral and neural responses to the odor.

**Figure 1:**
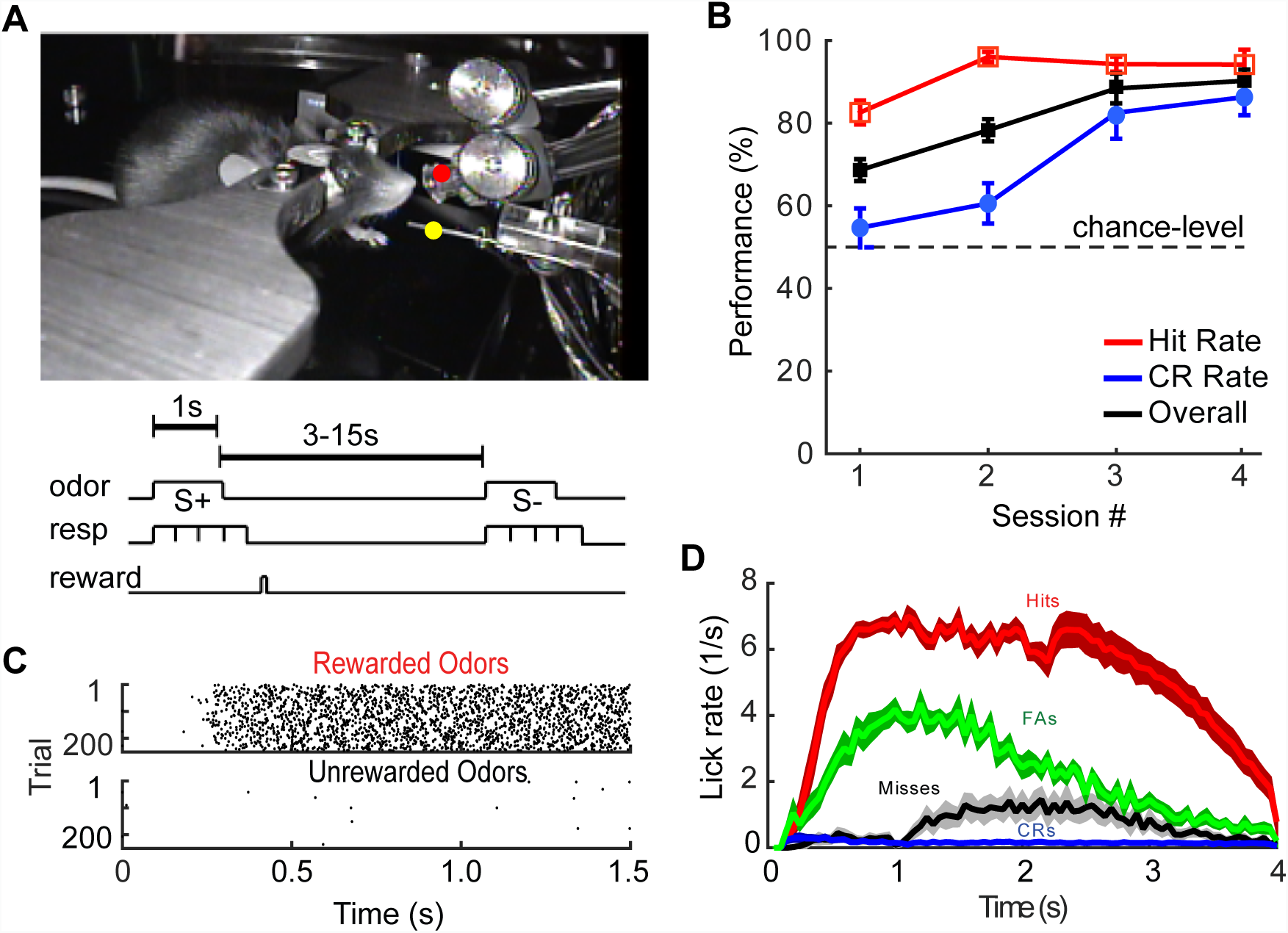
Behavioral task design and mouse performance. **A**. (top) Mouse in the head-fixed behavior setup. The odor port (red dot) and lick port (yellow dot) are labeled. (bottom) Diagram of the task structure, indicating the periods for odor, response (lick/no-lick), and reward delivery. **B**. Average task performance over the first four sessions for 14 mice. The hit rate, correct rejection (CR) rate, and overall performance rate are shown separately. **C**. Raster plots of individual lick times by trial, shown separately for rewarded and unrewarded odors for an expert mouse. **D**. Lick rate by trial type averaged over sessions in well-trained mice (i.e. after session 5). Shading indicates standard error of the mean. FA: false alarm; CR: correct rejection. For panels C and D, 0 marks onset of odor stimulus.

Mice acquired the task structure and reached a high level of performance (>90% correct trials) within three to four sessions from the start of training with odors (one session per day, consecutive days; Figure 1B-D; 14 mice). Among correct behavioral responses, the fraction of go responses on rewarded odor trials (i.e. hit rate) started higher and peaked earlier than the fraction of no-go responses on unrewarded odor trial (i.e. correct rejection rate), reflecting an initial behavioral bias towards licking. Upon further training, mice consistently began licking within 200-400ms of odor onset (Figure 1C) for rewarded odors and maintained licking until reward delivery but did not even initiate licking on most unrewarded odor trials (Figure 1C), indicating that the animals became confident of the reward association early during odor presentation. In a typical session, mice worked for about 300-500 trials and collected 150-250 water rewards before they were satiated.

Naive mice were initially trained on the task with the same panel of eight odors each day (one session per day) until they achieved successful performance (>90% correct) for two consecutive sessions. Following successful learning of the task structure, we replaced a subset of the odors with novel odors to study learning of odor-reward associations which were typically learned to near-saturating (>90% correct) performance within one or two subsequent sessions. This process was repeated in daily sessions that continued for one to two months in the same mouse. Thus, the rapid learning of novel odor-reward associations permitted the observation of behavior and physiology during the acquisition of many odor-reward pairs. We first present analysis from sessions with odor-reward pairs in which successful learning had already been demonstrated through saturated high behavioral performance in a previous session (“familiar odors”) to characterize odor and reward tuning properties in pPC and OT. Subsequently, we describe the changes in behavioral and neuronal responses during the sessions in which novel odor-reward pairs were introduced and learning took place (“novel odors”).

Since neural responses to odors is aligned to respiration (Cury and Uchida, 2010; Shusterman et al., 2011), and behaving rodents can alter the rate of sniffing significantly (Kepecs et al., 2007; Welker, 1964; Wesson et al., 2008), we monitored respiration with a cannula implanted over the nasal cavity, contralateral to the side where tetrode recordings were performed. Once familiar with the task structure, the rate of sniffing during odor presentation did not differ from the spontaneous rate between trials (3.8 ± 0.52 s^−1^ vs. 3.9 ± 0.45 s^−1^; Supp Figure 1).

### Task-related activity in pPC and OT

We used multi-tetrode drives (Gray et al., 1995) to isolate and record single units in either pPC (n=6 mice; 385 isolated units) or OT (n=8 mice; 270 isolated units). Following the completion of all behavior and recordings, tetrode placement was confirmed with an electrolytic lesion and postmortem histology (Supp Figure 2). Peri-stimulus time histograms (PSTHs) are shown for representative pPC and OT cells in Figure 2A,B. Since odor responses began following the first sniff during odor presentation, all of the remaining analyses were performed using the time of the first sniff as the start time for each trial. An example of sniff-aligned and odor onset-aligned activity for the same OT cell is shown in (Figure 2A) for comparison.

**Figure 2:**
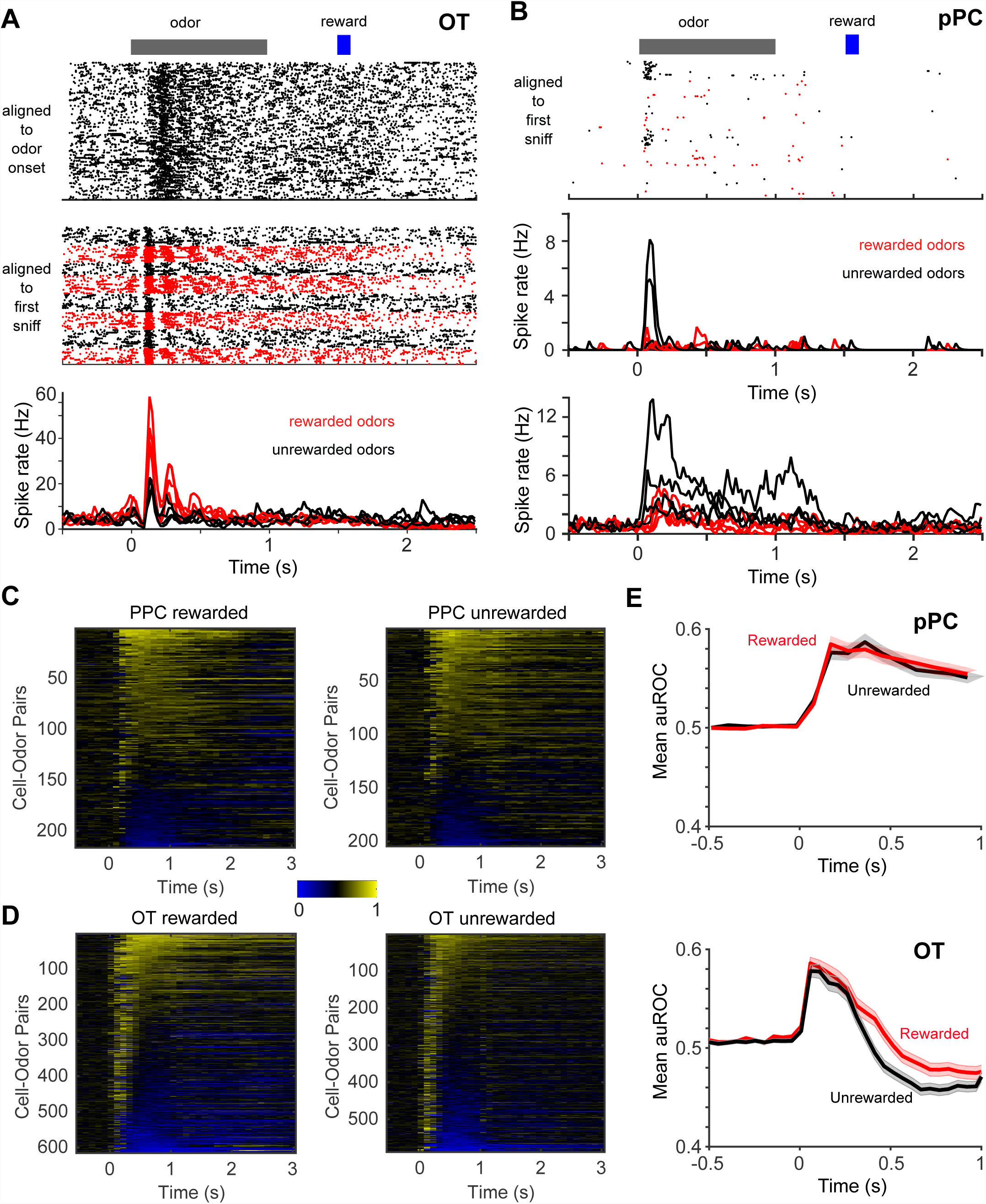
Responses of pPC and OT neurons. A. An example OT cell. Spike rasters are shown unaligned (top) and aligned (middle) to the first inhalation of odor. A peri-stimulus time histogram (PSTH) aligned to the first sniff is shown directly below the rasters. Four PSTHs for the rewarded odors are shown in red and four PSTHs for the unrewarded odors are shown in black. B. Spike rasters (top) and PSTHs (middle) for an exemplar cell from pPC. Bottom: another cell with more sustained odor responses. C, D. auROC timecourse for each cell-odor pair for cells recorded in pPC and OT separated into rewarded and unrewarded odors. Odor onset is at 0s. Yellow indicates that the cell was excited relative to its baseline firing rate, whereas blue indicates that the cell was inhibited relative to its baseline firing rate. For visualization purposes, cell-odor pairs with noisy firing during pre-odor time are not shown, resulting in a different number of cell-odor pairs between rewarded and unrewarded odors despite equal number of recorded cell-odor pairs.

To characterize the responses of neurons to odors, we used the metric of area under the receiver-operating characteristic curve (auROC) for each cell-odor pair relative to baseline (i.e. pre-stimulus) firing rate ((Green and Swets, 1966); see Methods). We plotted the responses for all significant cell-odor pairs (at least one 100ms bin different from baseline at a significance level of p < 0.01) in both areas with rewarded and unrewarded odors separated to look for differences in prevalence of specific type of activity (Figure 2C,D). With this criterion, 69 units in pPC and 177 units in OT responded to at least one odor. This significance measure was used only for visualization and initial characterization, and further analysis (such as decoding) used all recorded neurons.

Baseline firing of pPC neurons was low (2.33 ± 3.77 Hz, n = 372 units; Supp Figure 2) and odor responses were predominantly excitatory (Figure 2C). As expected, neurons with excitatory responses showed odor selectivity, responding only to a subset of odors (Figure 2A,B). Many cells continued to respond for the duration of the one second odor presentation, seen as sustained band of yellow (Figure 2C). Other cells had a strong transient (100-200ms) response well aligned to the first inhalation of an odor, but did not respond for the remainder of the odor presentation despite continued sniffing (Figure 2C). Interestingly, the cells that were inhibited appear to be inhibited by all odors non-selectively. Low baseline firing rates made it difficult to detect odor-selective inhibition, if it were present, and suggest that these neurons did not directly contribute to odor discrimination. Therefore, inhibitory responses of pPC neurons were not analyzed further.

The responses of OT neurons were clearly distinguishable from those of pPC neurons in several ways. Higher baseline firing rates (e.g. greater than 5 Hz) were much more common in OT (16.2 ± 16.9 Hz, n = 270 units; Supp Figure 2). Unlike pPC, a substantial portion of neurons were inhibited during odor presentation, often following a brief period of excitation (Figure 2D). Transient excitatory responses at the beginning of odor presentation were much more prevalent in the OT (Figure 2D). OT neurons tended to be less odor selective according to a lifetime sparseness measure (Willmore and Tolhurst, 2001), but neurons in both regions spanned a wide range of selectivity (Supp Figure 2).

### Reward-related activity at the single neuron level

We characterized reward coding in the pPC and OT, first focusing on familiar odors (i.e. odors that the mouse successfully learned in a previous session). Responses lasting longer than 500ms were more common in OT for rewarded odors than unrewarded odors (Figure 2D). In contrast, the overall distribution of response profiles did not differ between rewarded and unrewarded odors for pPC neurons (Figure 2C). This difference is clearly visible in the global average responses for rewarded vs. unrewarded odors for both areas, with OT neurons (but not pPC neurons) clearly showing divergent responses starting around 300 ms after onset of the first sniff (Figure 2E).

We then examined the extent to which reward information was explicitly available in the firing rate of individual neurons. Firing rates of individual neurons in each area suggested that OT neurons may be more discriminating between rewarded and unrewarded odors than pPC neurons (Figure 3A,B). An auROC analysis comparing responses to all rewarded odors to all unrewarded odors revealed that reward selectivity was substantially more prevalent in OT than pPC both during and after odor presentation (Figure 3C,D). Furthermore, roughly equal numbers of OT cells responded with higher firing rates for rewarded odors than unrewarded odors (yellow in Figure 3D) versus higher firing rates for unrewarded odors than rewarded odors (blue in Figure 3D). In fact, some cells switched preference (i.e. higher firing rate) between rewarded and unrewarded odors during the course of the response.

**Figure 3:**
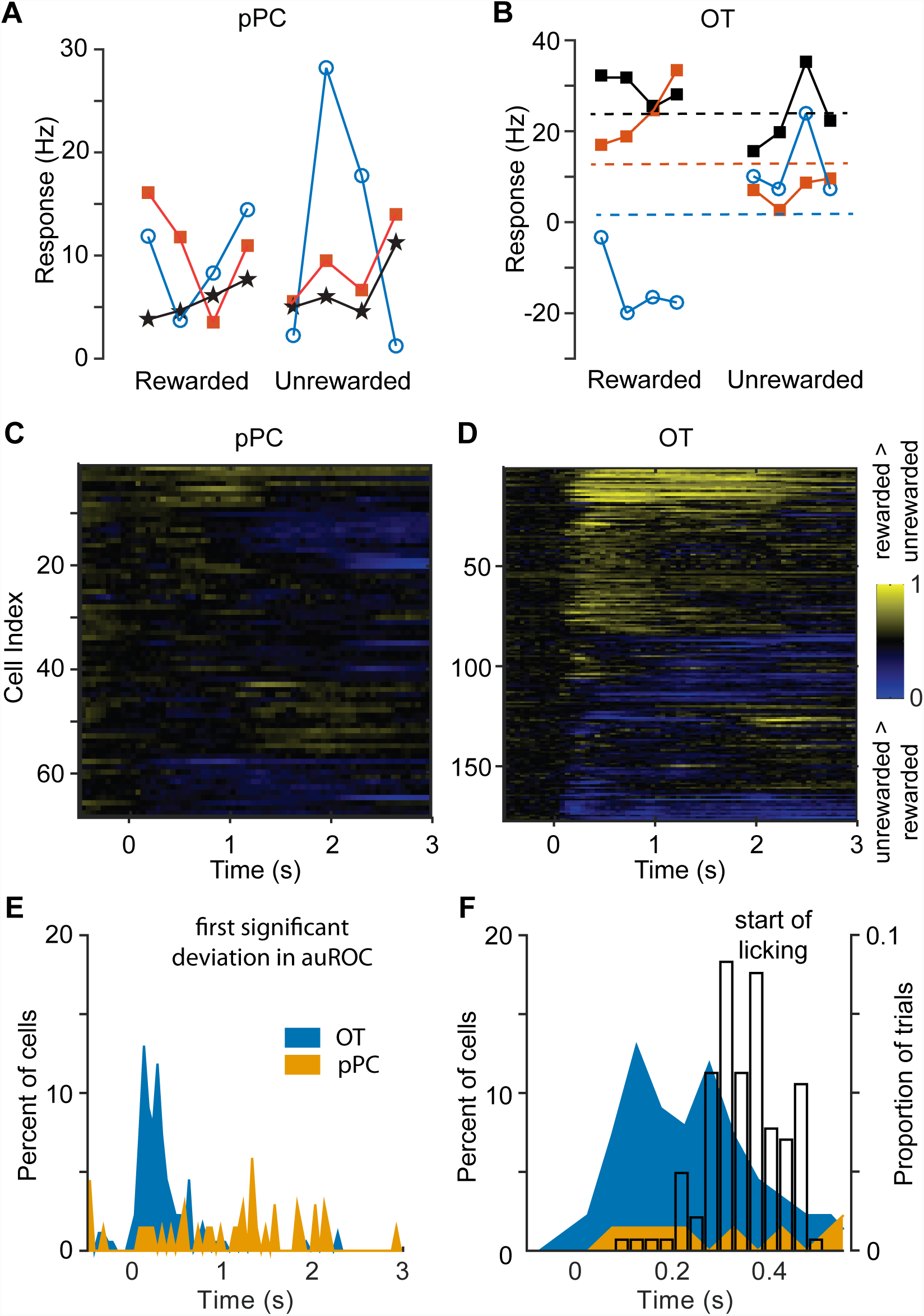
Reward valence selectivity during decision-making. **A**. Responses of three examplar cells from pPC to 8 odors, with no clear reward category information. B. Three exemplar cells from OT, two of which (red and blue) show clear reward selectivity (visualized by dashed lines separating firing rates), and one (black) showing imperfect reward selectivity. **C, D**. auROC to illustrate cells’ ability to discriminate rewarded versus unrewarded odors that had been successfully learned in previous sessions (i.e. familiar odors). Responses are aligned to the first inhalation following odor onset (t=0). The cells (rows) are hierarchically clustered by the first three principal components calculated from the initial (first 200ms) firing rate, delayed response rate (200ms-800ms), and response rate at the time of reward delivery (1500-2000ms). **E**. Histograms of the first time bin in which the auROC for a cell was significant for cells in pPC and OT. **F**. Zoom-in of the histograms in **E**, along with an overlay (hollow black bars) of the mean time of the first lick on hit trials to illustrate the beginning of the motor response.

We then asked whether reward selectivity emerged at times corresponding to particular task events, such as stimulus onset or reward delivery. In order to measure the significance of reward selectivity, we compared the auROC at times after odor onset against the distribution of auROCs during pre-stimulus time. Consistent with a lack of actual reward selectivity for pPC cells, the times at which the reward auROC first reached a very liberal threshold of significance (p < 0.01, 35 comparisons per cell for the 100ms time bins spanning the 3.5s window; family-wise type 1 error rate = 0.296) were scattered throughout the trial and not well-aligned to any task events (Figure 3E: orange). In contrast, reward selectivity emerged in 46% of OT cells within the first 300ms of odor sampling (Figure 3E: blue). The absence of any subsequent peaks indicates that there was not a separate population of OT neurons that responded only to later task events, such as reward delivery. The majority of these selective responses emerged before the mouse began to lick on rewarded odor trials (Figure 3F). This difference in reward selectivity between neurons in pPC and OT was not simply due to differences in stimulus selectivity and firing rates in the two regions (Supp Figure 3).

### Reward coding at the population level

Collectively, many individual cells with weak reward valence selectivity could still give rise to reliable valence selectivity at the population level. To determine whether our pPC and OT datasets had this property, we attempted to decode the valence of odors from single trial responses of cells in each area using a support vector machine with a linear kernel (Hastie et al., 2009). For each cell, the trials during the session it was recorded were randomly assigned to training (80%), validation (10%), and test (10%) sets.

To empirically determine an appropriate time window to analyze, we decoded reward valence from OT activity for a series of windows beginning at the first inhalation of the stimulus; specifically, we used the average firing rate response (i.e. baseline subtracted) for each cell within the window. Decoding performance improved with progressively longer time windows until saturation at ~200ms (Figure 4A). Accordingly, all subsequent decoding analyses were performed using the average firing rate response within a 200ms window beginning at the first sniff of the odor.

**Figure 4:**
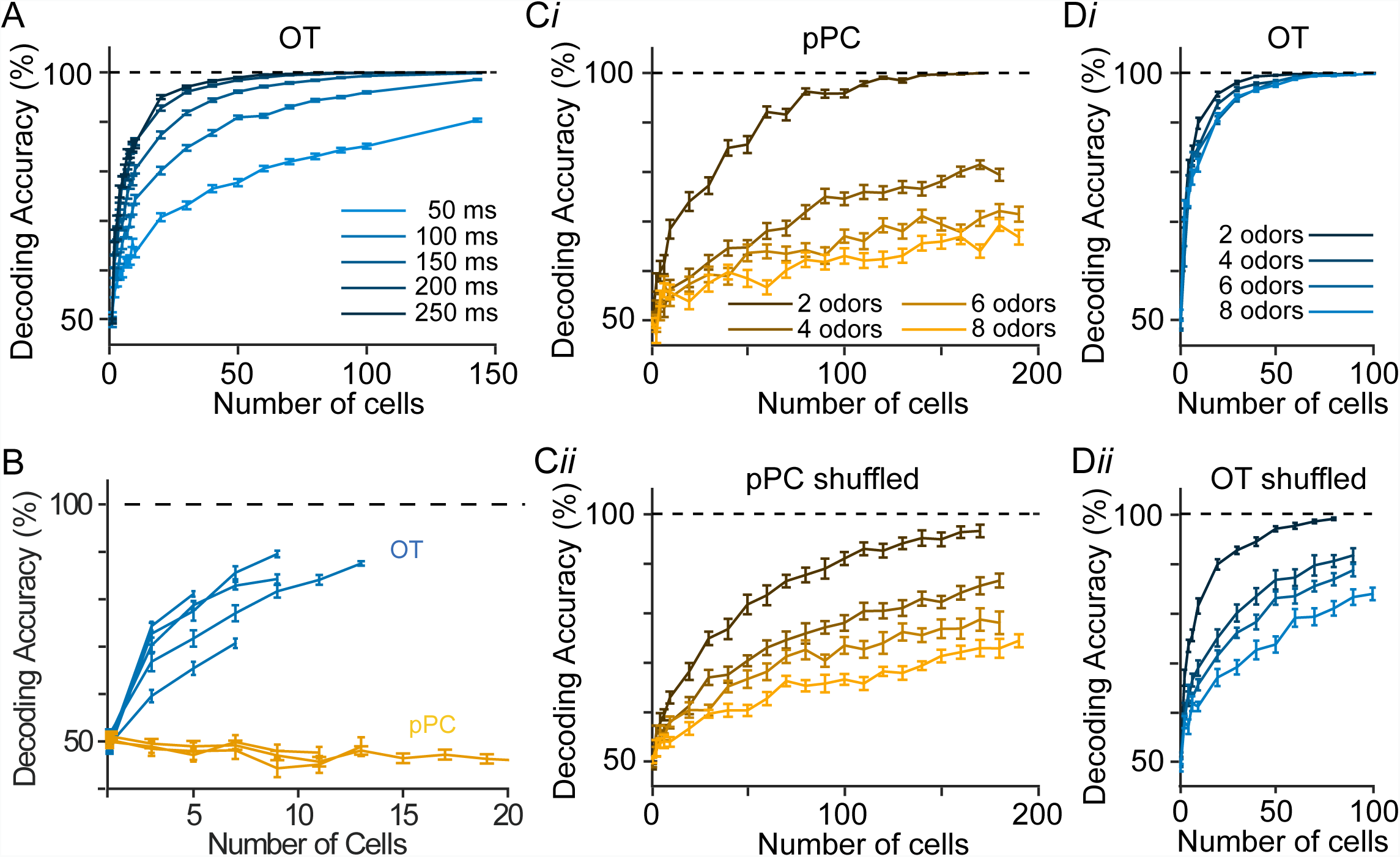
Decoding reward valence from population data. **A**. Decoding accuracy for reward valence obtained with a support vector machine, trained and tested on the average response rate of OT cells in a window that began at the time of the first inhalation of an odor and had a duration equal to the time denoted for each curve. **B.** Training and testing linear classifiers on different sets of odors. Support vector machines with a linear kernel were trained on the single-trial responses to one set of 4 odors and tested on a different set of 4 odors. Each line corresponds to the test performance for a different mouse or set of test odors. pPC (yellow): 3 curves total; each curve is a different mouse (each mouse had a different set of test odors). OT (blue): 5 curves total: 3 curves for three mice with the same test odor set, 1 curve for a second set of test odors for one of the three mice, and 1 curve for a fourth mouse with a different set of odors than the other three mice. **Ci, Di**. Decoding accuracy for similar classifiers that were trained and tested on the given number of odors (color correspondence indicated in the inset), half of which were rewarded odors and half of which were unrewarded odors. **Cii, Dii**. The same as Ci and Di but for classifiers trained and tested on odor responses with the reward valence label shuffled.

To directly test for an explicit population code for reward valence, we trained linear classifiers (SVMs) to decode reward from the responses to a set of four odors (2 rewarded, 2 unrewarded), then tested the ability of the same classifier to decode reward from the responses to the other set of four odors presented in the same session (Figure 4B). Overall, we found that classifiers trained on OT data successfully decoded reward from responses to odors that the classifiers did not see during training. On the other hand, classifiers trained on pPC data did not have this property. Importantly, these results were consistent across individual mice (OT: n=5 mice, pPC: n=3 mice) and odor sets for which populations of at least 5 cells were available (Figure 4B; Methods)

A single neuron could respond similarly to two odors with the same valence, because the response is valence-specific or strictly due to similar purely sensory-driven responses evoked by the two odors. In turn, the activity of a population of neurons might also falsely appear to discriminate rewarded versus unrewarded odors due to chance (i.e. the presence of sensory but not reward information) when probing with a small number of odors. If population activity explicitly codes for reward valence beyond sensory tuning alone, however, a single classifier should be able to decode the reward valence for increasingly larger number of odors without performance declining because the reward information persists despite the presence of extraneous stimulus-specific information. Therefore, we tested the ability of a decoder to classify reward valence from progressively larger number of odors (e.g. 1 rewarded odor versus 1 unrewarded odor, 2 rewarded odors versus 2 unrewarded odors, etc.). To minimize the contribution of stimulus tuning for any single odor to the overall ability to successfully categorize rewarded versus unrewarded odors, we leveraged the fact that the data collected from each mouse comprised many odor-reward associations, aggregated from individual sessions with 8 odors each. We therefore trained and tested the classifiers on this pooled dataset with balanced numbers of rewarded and unrewarded odors (14 mice; minimum of 5 sessions per mouse). Importantly, the size of the training set (i.e. number of single-trial responses) was held constant regardless of the number of odors used.

In this test, the decoder successfully classified pPC responses to two odors (one rewarded and one unrewarded), but performance declined rapidly when challenged to decode valence from responses to larger numbers of odors suggesting that pPC odor responses conveyed little to no linearly-decodable information about valence (Figure 4Ci). As a control, we compared this result to the decoding performance achieved by a classifier that was trained and tested on the same dataset after shuffling the valence labels of the odors. This shuffling procedure would remove any potential explicit reward information and force the decoder to rely on stimulus-tuning alone for classification. Decoding performance on valence-shuffled pPC data followed the same trend as the unshuffled pPC data (Figure 4Cii), again suggesting the absence of an explicit code for reward in pPC. In contrast, decoder performance for the OT data did not decrease substantially as the number of odors increased (Figure 4Di), but did decrease for valence-shuffled OT data, confirming that valence-specific information was lost in the shuffling process (Figure 4Dii).

These results clearly indicate that the reward category of odors could be explicitly read out from small groups of OT neurons, but not from pPC neurons.

### Multiplexing of odor and reward codes

To assess the degree to which odor identity information is represented in the activity of pPC or OT neurons, we picked sets of four odors (two rewarded and two unrewarded) for which we had the most cells recorded during sessions in which all four odors were used (mean: 24 cells, range: 12-45 cells). We quantified odor identity information as the decoding accuracy when classifying responses to one odor versus responses to the other three odors (i.e. “one versus rest” classification). For each brain region, we tested separately the performance of different mice and sets of odors. In OT, decoding of any individual odor was significantly better than chance (Figure 5A). Decoding of individual odors was much worse in pPC than OT, close to chance (Figure 5A). This is likely due to sparse odor responses in pPC (Supp Figure 2) resulting in few or no responses to a given odor within the relatively small populations of cells available for this analysis (mean: 32 cells per odor set in pPC). This analysis indicated that neurons in OT carry information about odor identity multiplexed with reward information.

**Figure 5:**
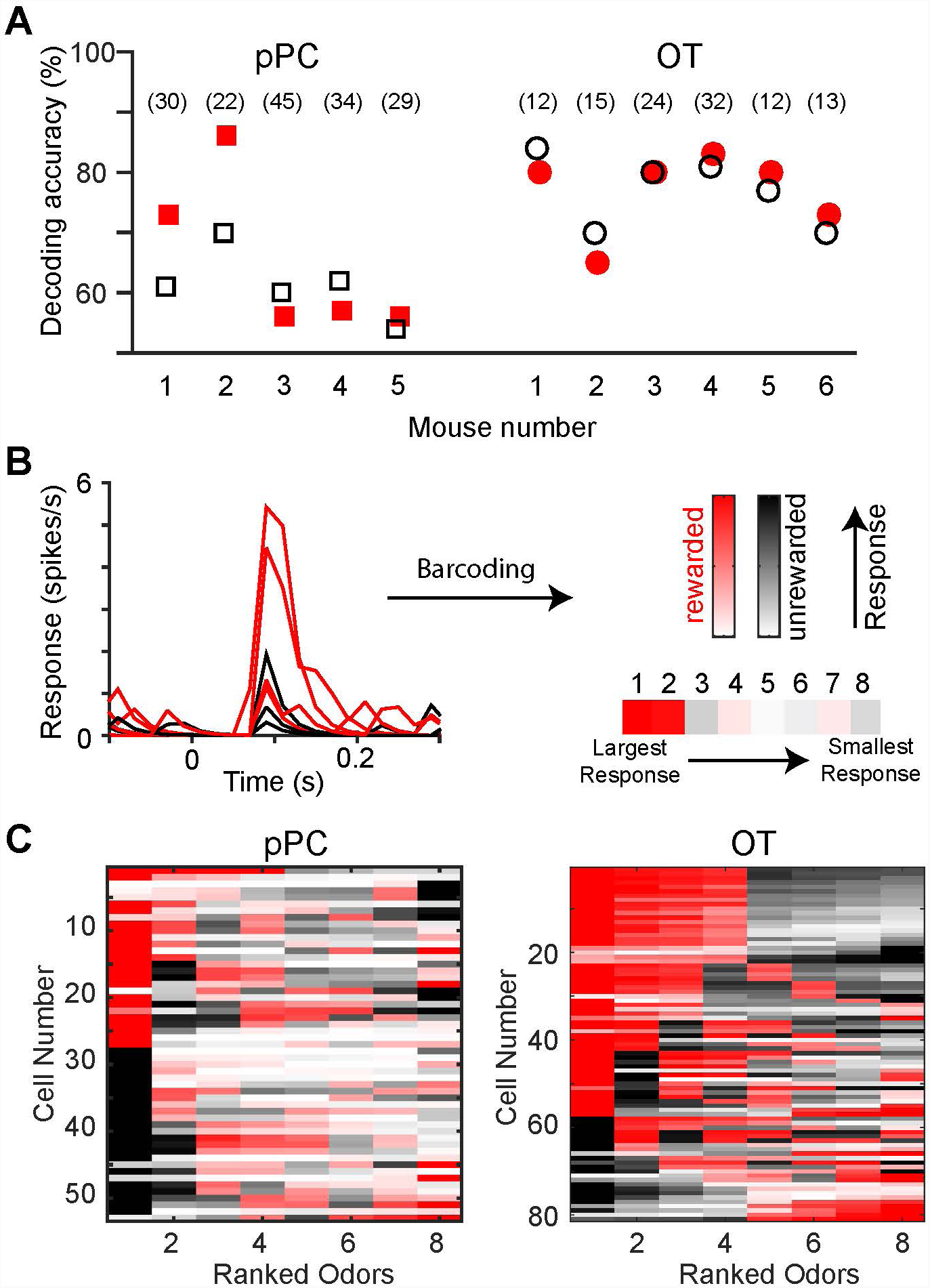
Multiplexing of odor and reward codes. **A.** Decoding accuracy for unrewarded (black) and rewarded (red) odors are shown (each point is average of two odors in each category). Data for each mouse for each region shown individually, with number of cells in brackets at top. Chance performance is 50% for all classifiers. **B.** Single-cell peri-stimulus time histograms are used to produce barcodes that illustrate both odorand reward sensitivity of a cell’s response during the first 200ms of odor presentation, beginning with the first inhalation. Odors are rank ordered from most positive-going response to most negative-going response, then colored according to valence (red - rewarded, black - unrewarded) and the normalized response to the odor. **C.** Stacks of barcodes for cells recorded in pPC and OT during sessions in which all eight odors in the stimulus panel had been successfully learned in a previous session (i.e. familiar odors).

To visualize the multiplexing of odor and reward selectivity, we created a bar code for each neuron that separates responses to each of the eight individual odors but also distinguishes odors based on their reward valence (Figure 5B). The position of each odor in the bar code corresponds to the rank-order of the response magnitude (during the first 200ms window), from most positive on the left to most negative on the right. The color of each segment corresponds to the valence of the odor (red=rewarded; black=unrewarded) and the intensity of the color corresponds to the odor-evoked response normalized to the strongest response among the eight odors. The combined barcodes from neurons in the two regions illustrate that pPC neurons have sparse selectivity for odors and do not discriminate valence, whereas OT neurons respond much more densely to odors and partially, but not entirely, discriminate valence (Figure 5C).

### Novel odor learning

The results described thus far established that OT neurons had an explicit representation for reward valence that was multiplexed with odor tuning, whereas pPC neurons lacked an explicit representation for reward valence. Next, we investigated the evolution of reward coding during the process of learning novel odor-reward associations, which we were able to do within recording sessions. Approximately every other session following successful learning of the task structure (total of 71 sessions), four of the eight odors that had been used in previous sessions were replaced with four novel odors (two rewarded and two unrewarded) (Supp Figure 4A). Mice were not exposed to the odors prior to training and reward valence was randomly assigned to each odor for every mouse.

We found that across different mice and different novel odors (8 mice, 71 sessions) rapid learning occurs in the very first session, with additional improvement in performance over the next two sessions (Figure 6A,B). Correct go responses to rewarded odors become frequent earlier than correct no-go responses to unrewarded odors (Supp Figure 4). High accuracy is maintained in these sessions for the already familiar odors (Figure 6B, right).

**Figure 6:**
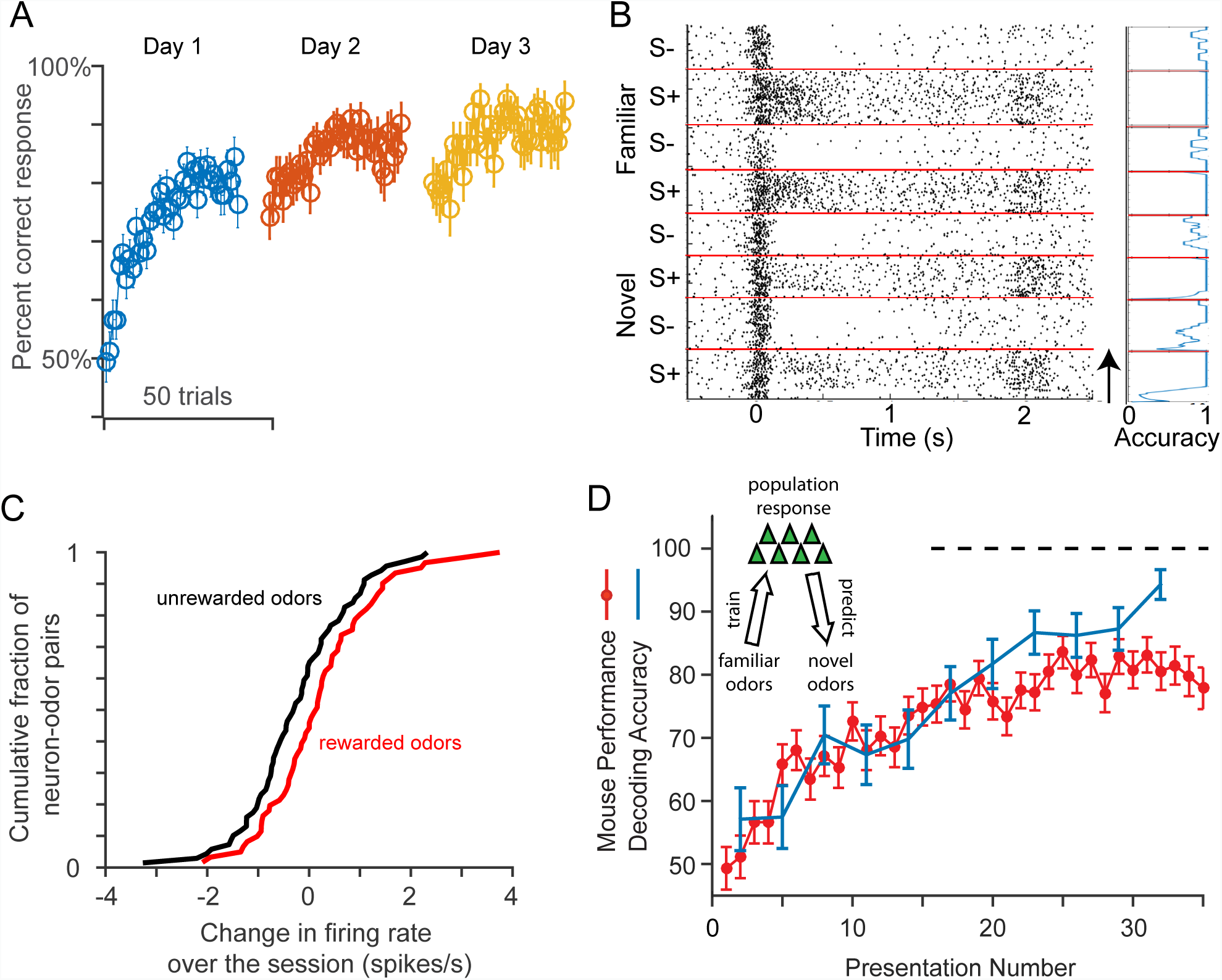
Novel odor learning. **A**. After 5-10 sessions of training with 8 familiar odors, half of the odors (2 rewarded and 2 unrewarded) were replaced between some sessions. The performance for the 4 novel odors on the first 3 days of learning is shown averaged for all novel odors as a function of presentation number. Each novel odor was presented ~35 times (randomly) in each session. **B**. Responses of an example neuron to the 4 rewarded (S+) and 4 unrewarded (S-) odors, half of each were familiar and novel. Trials were randomly presented, but are shown sorted for visualization. Within each category, trials are displayed in order of presentation in the session, with early trials at bottom (arrow at bottom right). The accuracy of performance for each of the odors is indicated at right. Note that accuracy for familiar odors is near 1 for all trials, but for novel odors accuracy starts low and approaches 1. **C**. Based on the average firing rate (in 200 ms after first sniff) for each neuron-odor pair, the difference in firing rate between the last 10 and first 10 trials was calculated for novel rewarded and unrewarded odors. As a population, neuronal responses for rewarded odors become slightly (p < 0.05) stronger compared to unrewarded odors. Each of the recorded neurons was only recorded during a single session, and therefore had a different stimulus panel than neurons recorded during other sessions. **D**. Decoding reward selectivity from OT responses to novel odors. *Inset:* A classifier was trained to discriminate two familiar rewarded odors from two familiar unrewarded odors, then tested on its ability to predict reward valence from responses to two novel rewarded and two novel unrewarded odors. Decoding accuracy (blue; averaged across 71 novel sessions) steadily increased with repeated presentation of novel odors, and tracked performance of mice (red). Error bars show standard error of the mean.

We recorded activity of OT neurons during novel odor learning, with four familiar and four unfamiliar odors presented in randomly interleaved trials (Figure 6B). We asked whether the activity of individual neurons changed during the course of the session by examining the difference in firing rate for the first and last 10 trials of an odor. For each OT cell recorded during learning, we calculated the change in response to each novel odor normalized to the mean and standard deviation of the cell’s responses to the four familiar odors used in the same session. Across the population of neurons, we found that the distribution of these normalized response changes for rewarded novel odors was significantly different (mean change of 0.22 +/-0.13 spikes/s, 69 cells, p = 0.02) from the distribution for unrewarded novel odors (mean change of - 0.22 +/- 0.14 spikes/s, 61 cells, Figure 6C).

To determine whether odor-reward associations were reflected in the activity of populations of OT neurons during learning, we leveraged the fact that only four of the odors were novel in learning sessions and the other four odors were familiar (i.e. learned in a previous session). We therefore compared the evolution of reward selectivity of novel odors using the familiar odors as a reference point (Figure 6D, inset). To this end, we trained a linear decoder to classify single-trial responses (mean firing rate response in first 200ms of odor sampling) to *familiar* rewarded versus unrewarded odors, then tested the performance on *novel* rewarded and unrewarded odors. First, we quantified the density of reward valence coding in OT by determining the minimum population size for which this training procedure resulted in decoding accuracy that approximated the average behavioral performance of a well-trained mouse (~90%) on familiar odors alone – this size was 20 cells (Figure 4C). The average performance of 100 classifiers each trained on a random group of 20 cells out of the 73 recorded during single-session learning is shown in Figure 6D. Indeed, there is a highly significant (p<10^−6^) increase in decoder performance as a function of the presentation number (i.e. trials with the same odor), starting near chance performance at the beginning of the session and reaching comparable performance to familiar odors (>90%) by the end of the very first session with the novel odors. Remarkably, the time course of the improvement in decoding closely tracks the performance of mice in discovering the valence of the novel odors (Figure 6D). Overall, these results establish that reward selectivity in the OT develops in parallel with learning at the behavioral level.

### Relating neural activity and behavior

If the activity in OT is used to drive behavior, we would expect a stronger relationship between coding and behavior (i.e. “Consistency”) (Majaj et al., 2015) in OT than other candidate areas, such as pPC. Indeed, the relationship between the decoding accuracy for individual odors and behavioral performance of the mouse for the corresponding odors was stronger in OT than pPC (r = 0.40 in OT, p = 0.05; r = 0.19 in pPC, p > 0.2; Figure 7A,B).

**Figure 7:**
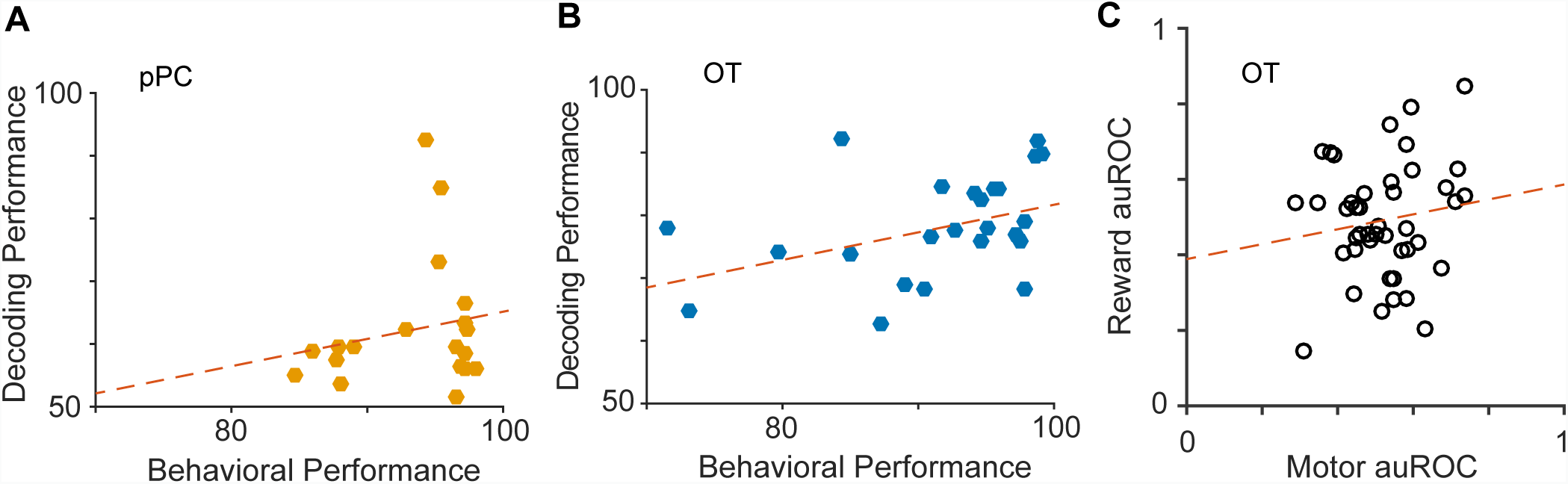
**A.** Odor-by-odor relationship between behavioral performance and neural coding for pPC. Each dot is a single odor. N = 19, r = 0.19, p = 0.41. **B**. Same plot as in A, for OT. N = 24, r = 0.4, p = 0.05. **C**. OT activity selectivity on error trials. The effect of motor (lick/no-lick) decision was measured as the auROC when discriminating responses during the first 200ms of false alarm versus correct rejection trials. Similarly, the effect of reward valence was measured as the auROC when discriminating hit versus false alarm trials. A minimum of 15 false alarm trials was required for the cell to be included in the analysis. N = 43, r = 0.14, p = 0.36.

Finally, we examined whether OT responses were more closely related to reward valence or motor decision. The go/no-go task structure ties reward valence to motor decision (i.e. go=rewarded, no-go=unrewarded), so we examined OT responses on error trials in order to dissociate these variables. Due to the asymmetry in hit rate versus correct rejection rate in our task, there were too few misses to be used for this analysis; instead, we focused on false alarm trials. We quantified the reward valence coding by calculating the auROC of OT neuron responses across hit versus false alarm trials for which the motor decision was the same but the valence differed (“reward auROC”). Likewise, we quantified the motor decision coding by calculating the auROC of OT responses across correct rejection versus false alarm trials (“motor auROC”), for which the reward valence was the same but the motor decision differed. Importantly, we never included responses from the same odor among both the false alarm trials and correct rejection trials when calculating the motor auROC. This ensured that we did not artificially decrease the motor auROC by forcing the discrimination of an odor from itself, relative to the valence auROC in which the odors on hit versus false alarm trials were always different, by construction. If motor decision was the main driver of selective responses in OT neurons, there should be a strong correlation between motor auROC and valence auROC across a population. Instead, we found that these two measures were not significantly correlated (p = 0.36) across the population of OT neurons (Figure 7C), indicating that the observed reward tuning is not primarily driven by a difference in motor behavior.

## Discussion

In this work, we have shown that mice learn to successfully perform an odor-reward categorization task over the course of a few days. Once the task structure had been learned, mice could flexibly apply the task rules in order to acquire novel odor-reward associations within a single session. This categorization task allowed us to study how odor and reward coding interact in the olfactory system.

We observed a code in pPC that had sparse odor responses and weak reward selectivity and a code in OT that had strong reward selectivity and less sparse odor responses than pPC. Reward selectivity in OT emerged in the pre-decision time during the first sniff of odor cues and was detectable, though weak, at the single-cell level. Moreover, reward selectivity and successful behavioral performance developed in parallel as novel odor-reward associations were learned. These results provide important constraints on how odors are processed in the olfactory system during reward-motivated behaviors.

### Identity coding

Although much previous work has focused on coding of odor *identity* in the piriform cortex (Bolding and Franks, 2017; Gottfried, 2010; Illig and Haberly, 2003; Iurilli and Datta, 2017; Roland et al., 2017; Stettler and Axel, 2009; Wilson and Sullivan, 2011), our experiments were designed to focus on reward-related activity and only a small number of neurons were recorded during experimental sessions with a particular set of odors. Previous studies (Bolding and Franks, 2017; Iurilli and Datta, 2017) and our own unpublished work (Grimaud et al, in preparation) demonstrate that more than 100 cells are necessary for odor identity decoding in the piriform cortex. Indeed, identity decoding was very poor in pPC in our current study, likely due to the sparse odor responses in pPC resulting in few or no responses to a given odor within the relatively small populations of cells available for this analysis (mean: 32 cells per odor set in pPC). Interestingly, OT neurons had denser responses than pPC neurons, which allowed easier decoding of odor identity, but individual OT neurons did not typically respond to all rewarded or unrewarded odors. Therefore, individual OT neurons carry some information about both odor identity and reward category. This multiplexing of information in OT may allow rapid and flexible learning of odor valence.

### Reward coding

A major finding of our study is that neurons in the OT carry an explicit representation of reward category, independent of the identity of the odor. This information can be gleaned reliably from the firing rates of only a few dozen arbitrarily selected neurons, within a brief period (~200 ms) after odor sampling. Remarkably, when mice learn new odor-reward associations, neurons in the OT alter their firing properties to begin coding for reward category in concert with learning during the same session. Importantly, we show that the activity we find to be correlated with reward valence is not simply a consequence of the motor action. First, the emergence of reward information in the activity of OT neurons precedes the motor action (licking). Second, when using error trials, the ability to decode the choice the mouse makes (licking or not licking) for each neuron is unrelated to the ability to decode the reward category. Therefore, while OT neurons may also carry information related to the motor action, this would be in addition to the information they carry about reward category.

There is significant anatomical and pharmacological work implicating the OT in reward and motivated behavior (Heimer et al., 1982; Ikemoto, 2003, 2007). Recent work has implicated the OT in valence coding (Gadziola and Wesson, 2016; Gadziola et al., 2015; Zhang et al., 2017b) though the relationship between valence coding and stimulus (odor) tuning was not clear. Here, we used a large number of odors to probe the breadth of stimulus responses (i.e. lifetime sparseness) when many different odors reliably predict reward or no reward. We find that OT neurons multiplex odor identity and reward category information. In addition, we find that the mouse’s behavioral performance for an individual odor is correlated with the ability to decode reward category from OT activity in response to that odor. Monitoring sniffing in all the experiments allowed us to determine the timing of neural responses at high temporal resolution and revealed that reward category-related activity in the OT arises rapidly (on average, 100ms after onset of sniffing, more than 200ms before the average onset of the motor act). In contrast, we found little evidence for an explicit representation of reward category information in the pPC, beyond that which is expected merely from sensory tuning. This appears to be at odds with an earlier study (Calu et al., 2007) that suggested that the activity of pPC neurons can be modulated by reward. However, the study only used a few odors and a very small number of cells (less than 5) showed the effect, leaving open the possibility that the responses they observed were due to sensory tuning alone (as demonstrated in our Figure 4) or reflected later-onset brain-wide reward consumption anticipatory signals (Schultz, 2000). Together, these results suggest that the OT, not the pPC, is better situated to integrate olfactory and reward information to facilitate sensorimotor learning in our task.

### How do OT neurons acquire reward-selective responses so early during learning?

The emergence of reward-selective responses in a part of the striatum, such as the OT, is not unprecedented. Monkeys trained to associate reward with visual stimuli develop reward selectivity in the dorsal striatum that is apparent in the activity of individual neurons within a few trials of successful learning (Pasupathy and Miller, 2005; Schultz et al., 2003). In contrast, neurons in the prefrontal cortex took much longer to acquire reward selectivity (Pasupathy and Miller, 2005). OT, at least in rodents, has been suggested to implement similar computations for olfactory information that the rest of striatum implements for other modalities (Giessel and Datta, 2014; Wesson and Wilson, 2011). Indeed, OT and dorsal striatum have a similar composition of cells (e.g. ~98% D1- or D2-receptor-expressing medium spiny neurons), comparable density of input from VTA, and the same output targets (Wesson and Wilson, 2011). Our results strengthen the analogy between OT and dorsal striatum, adding physiological data to the known anatomical and cytoarchitecture similarities.

Reward-selective responses might emerge in striatum early during learning in order for plasticity in the striatum to support decision-making. Indeed, corticostriatal inputs have been shown to drive decision-making in an auditory discrimination task (Znamenskiy and Zador, 2013), and corticostriatal synapses exhibit plasticity over the course of training (Xiong et al., 2015). Moreover, it has been shown that the strength of olfactory synaptic inputs to the ventral striatum can be modulated by optogenetic release of dopamine in slices (Wieland et al., 2015), and the activation of VTA projections to the OT can induce odor preference in mice (Zhang et al., 2017b).

A recent study was published during the preparation of this manuscript, which arrives at some of the same conclusions as our study regarding the relationship of pPC and OT to coding odor-reward associations (Gadziola et al., 2019). Specifically, those authors also find and characterize the much denser reward coding in OT than pPC. Their approach involves a task design that is complementary to the one we have taken here, in which they use reward contingency reversals to demonstrate that odor-evoked responses in OT change in response to experimental manipulation of the reward valence predicted by the odor. On the other hand, our approach of training each mouse to learn many odor-reward associations enabled the dissociable characterization of multiplexed odor and reward valence components of the cells’ tuning in the OT. Taken together, along with the literature discussed already, our two studies lend strong support for a central role of the OT in odor-driven reward-motivated behaviors.

Our study sheds light on the representation of a behaviorally relevant aspect of the environment, reward, and how it relates to sensory coding in the olfactory system. The rapidity of odor-guided learning in mice, and the direct connectivity of the OT to the sensory periphery as well as motor and reward areas, make this behavior and brain region attractive candidates for the dissection of rapid sensorimotor reinforcement.

## Acknowledgements

This work was supported, in part, by a grant from the NIH (R01 DC017311) to VNM. DJM was supported by the National Science Foundation Graduate Research Fellowship under Grant No. (DGE1144152) and NRSA (F31 DC014602) from the NIDCD. Microscopy on fixed tissue samples was performed at the Harvard Center for Biological Imaging. We thank Nuné Martiros and Kenneth Blum for critical feedback that significantly improved the manuscript.

## Methods

### Animals (general)

All experimental animals were C57BL/6J mice obtained from the Charles River Laboratories, aged two to four months at the start of the experiments. Following surgical implantation of a tetrode drive and a custom head plate, all mice we rehoused individually. Experiments were conducted in accordance with Harvard University Animal Care Guidelines.

### Surgery

Surgeries were performed on naive animals and all behavioral training began after recovery from surgery. Mice were anesthetized with an IP injection of a mixture of Xylazine (10 mg/kg) and Ketamine (80 mg/kg). To access sniffing information, airflow was measured through a cannula implanted in the nasal canal. A craniotomy (~1mm) was made over the nasal canal (2mm anterior of nasal/frontal fissure, 1 mm lateral) on the skull, with a goal of implanting a nasal cannula to monitor sniffing during behavioral and physiological experiments. An 18 gauge stainless steel cannula (~5mm in length) was inserted into the craniotomy. To affix the cannula to the skull, superglue was used for initial placement followed by two applications of dental cement to ensure stability of the cannula. Additionally, a craniotomy (~1mm) was made over the dorsal skull at a location directly above the areas targeted for electrophysiological recordings with the goal of implanting a tetrode bundle (6 tetrodes plus a 200um-diameter optic fiber to ensure stability). The target locations were: pPC (coordinates: 0.5mm posterior, 3.8mm lateral from bregma, and 3.8mm ventral from brain surface) and OT (coordinates: 1.2mm anterior, 1.5-2mm lateral from bregma, and 4.6mm ventral from brain surface). To ensure stability of the animal’s head during behavior and recording at a later stage, a custom-made head plate (made of light-weight titanium, dimensions of 30 x 10 x 1 mm and weight of 0.8 g) was affixed to the skull. A shallow well was drilled over the posterior lateral skull and a single skull screw was affixed at that location. A wire was attached to this skull for grounding electrophysiological recordings. At the end of the surgery, a removable plastic cap was placed on the top of the nasal cannula to prevent foreign objects from entering the cannula. In addition, a plastic cone was positioned around the tetrode drive, and capped with a removable lid, to prevent damage to the drive. The nasal cannula and tetrode drive were implanted during the same surgery, but always on opposite (i.e. contralateral) sides. Each lateral olfactory tract delivers sensory input from a single olfactory bulb to the ipsilateral pPC and OT. Implanting the nasal cannula and tetrode drive on opposite sides was intended to avoid altering the natural airflow to the relevant sensory neurons for our recordings. Following the completion of the surgery, mice were given one week to recover.

### Odor stimulus delivery

A custom-made 16-odor olfactometer was used to deliver stimuli during behavioral tasks. The olfactometer had a carrier stream of air calibrated to flow at 1 liter per minute. A single odor at a time could be added onto this carrier stream by opening an odor-specific valve to permit airflow (at 0.1 liters per minute) from an input manifold, through a tube containing liquid-phase odorant, and finally into the carrier stream. One-way check valves prevented flow of odor from odorant-containing tubes into the carrier stream or back into the input manifold when air was not actively being flowed. Odor delivery to the animal was gated by a single (“final”) valve that directed odor to the animal or, between odor presentations, to an exhaust system. Note that this single valve was common to all odors and thus was not informative about odor identity. When odor was not flowed to the animal, a stream of clean air of the same flow-rate was directed to the animal. This ensured active clearing of odors between trials. In addition, an exhaust system cleared odors from the behavior chamber. To ensure immediate delivery of odors, and comparable odor concentration from trial-to-trial, the line to the final valve was primed with odor from the current trial beginning at the end of the previous trial (i.e. minimum of 5 seconds, sufficient to replace the line volume 3-5 times). Sound from the switching of an odor-specific valve to perform selection of an odor for the upcoming trial was masked by a substantially louder non-odor-specific valve turned on simultaneously. The valves corresponding to each odor were randomly switched occasionally between sessions to prevent animals from learning sound, rather than odor, cues. Finally, all tubing that was odor-specific was replaced at the same time as new odors were added to the olfactometer in order to prevent odor from previous sessions lingering and being used as cues for task performance.

### Behavioral task

When animals fully recovered from surgery (approximately 7 days), they were placed on a water deprivation schedule for behavioral conditioning. Animals were first acclimated to head restraint (with their implanted head plate held in place) on the behavior rig over the course of 3-5 days. Then, animals were trained to lick for water from a spout positions 2-3 millimeters from the mouth until satiated (typically 1-1.5mL); licking this spout (or not licking) would subsequently serve as the response for all behavioral tasks (go/no-go). Lick training took approximately 2-4 additional days. Next, odors were presented for one second with a varying (3 seconds minimum plus an exponential distribution with mean of 5 additional seconds) inter-stimulus interval, and reward was only available for a subset (half) of these odors. At first, only two odors (one rewarded, one unrewarded) were presented in order to expedite learning. The number of odors was increased to four and then eight (i.e. full task odor panel). Initially, water reward would be available immediately upon licking during odor presentation. Gradually, the delay between lick/odor presentation and reward delivery was increased until reward delivery occurred at least 500 milliseconds after the conclusion of odor presentation. To discourage a high false alarm rate, a hit was only scored on trials that the mouse licked in at least three out of four 350ms bins, beginning at the time of odor onset. It follows that mice were required to make a decision before the end of the second bin, 700ms into the odor presentation, but this did not seem to force early inaccurate decisions.

Neural recordings began once the animals reach strong performance (>90%correct) on the full eight-odor panel. No odors were replaced during this initial period in order to facilitate the acquisition of task structure. In the final stage of experiments, a subset of odors was replaced between some sessions in order to study learning of novel odor-reward associations.

### Respiration monitoring

Respiration was recorded through a nasal cannula implanted during surgery. Following surgery, the nasal cannula was cleaned daily to prevent clogging. In this way, nasal cannulae typically stayed clear continuously for months of recordings. During experiments, a plastic tube connected to a pressure sensor was fitted over the nasal cannula to form a continuous pressure environment. Pressure signals were amplified 10X, bandpass filtered 0.1-100Hz, and recorded at 1000 samples per second. For analysis, the start of each inhalation was identified as negative-going zero crossings in the pressure signal. Likewise, the end of each inhalation (i.e. start of exhalation) was identified as a subsequent positive-going zero crossing in the pressure signal.

### Electrophysiology

Neural activity was recorded with drives containing six tetrodes (Gray et al., 1995). The tip of each wire was gold-plated until the impedance was 250-450 kOhms. Electrophysiological signals were acquired and amplified with a custom-made system built on two 16-channel analog chips designed by Intan Technologies. Each channel was digitized, sampled at 20kHz, and bandpass filtered with a 2nd-order Butterworth filter 500-3000Hz. Potential spike events were detected as any activity that crossed 3.7 standard deviations of noise (corresponding to 1:10,000 events by chance). Potential spike events were then manually clustered with MClust software (David Redish) to identify single-unit activity. The quality of clustering was ensured by requiring less than 1 in 1000 spikes to have occurred within a 2 ms refractory period of one another, clear overlap of single spike waveforms, and L-distance below 0.05. Following the completion of all behavior and recordings, tetrode placement was confirmed with an electrolytic lesion and postmortem histology. To minimize selection bias, all spikes that were visible during each session were recorded, sorted and analyzed. At the end of the recording session each day, the entire bundle of tetrodes was lowered 40um to obtain a new set of neurons for the subsequent day.

### AuROC analysis

To characterize the responses of neurons to odors, we used the metric of area under the receiver-operating characteristic curve (auROC) for each cell-odor pair relative to baseline (i.e. pre-stimulus) firing rate. The auROC gives a measure of how well the firing rate at any given time can be discriminated from the baseline firing rate for that cell, independent of the absolute firing rates. Briefly, the value of the auROC for a given time bin indicates the percentile of that time bin in the distribution of baseline firing rates for bins of the same width. Therefore, a firing rate that is exactly the median of the baseline firing rate distribution would have an auROC of 0.5. Excitatory responses (yellow) correspond to an auROC greater than 0.5 but less than or equal to one, whereas inhibitory responses (blue) correspond to an auROC less than 0.5 but greater than or equal to zero. For analysis as well as data visualization, the auROC provides a more consistent measure to compare across neurons than absolute firing rate.

### Decoding

Decoding analysis (both identity and reward category) was done with SVMs with a linear kernel, using the standard 80% training, 10% validation and 10% test. Random shuffles from the entire population of neurons in a given region allowed many combinations, and therefore error estimates for decoding accuracy. Responses were obtained in a standard window of 200ms from onset of odor and first sniff inhalation (except in Figure 4A, where this window was systematically varied). Shuffle controls for decoding reward category were obtained by scrambling the valence assignment across the odors.

---

**Supplementary Figure 1:**
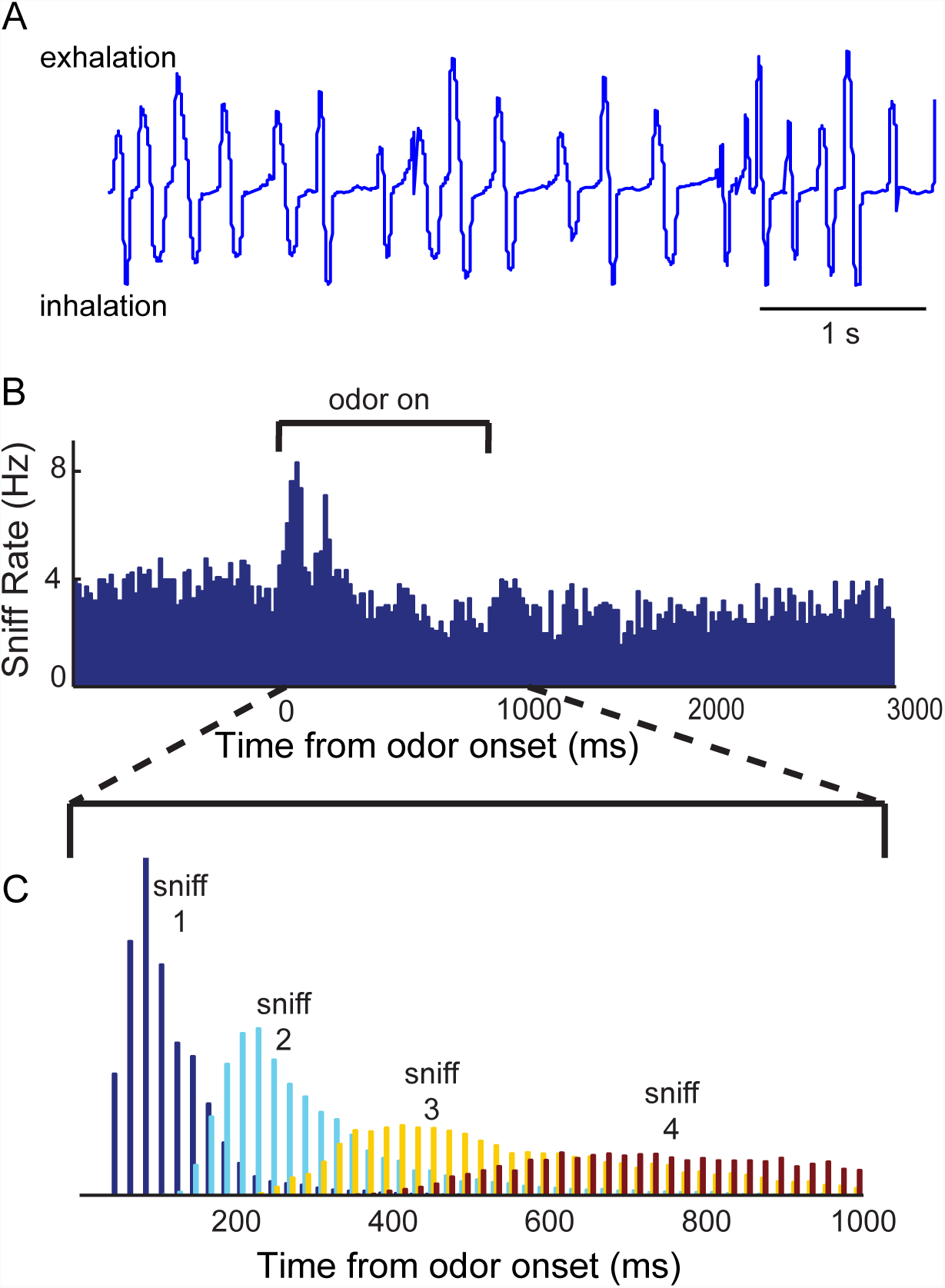
Sniff behavior during task performance. **A**. An example trace of five seconds of respiration measurements obtained through air pressure from a nasal cannula. Exhalation is positive-going pressure and inhalation is negative-going pressure. **B.** Average sniff rate over all trials from an example behavior session. **C**. Histograms of the times of the first, second, third, and fourth sniffs of each trial.

**Supplementary Figure 2:**
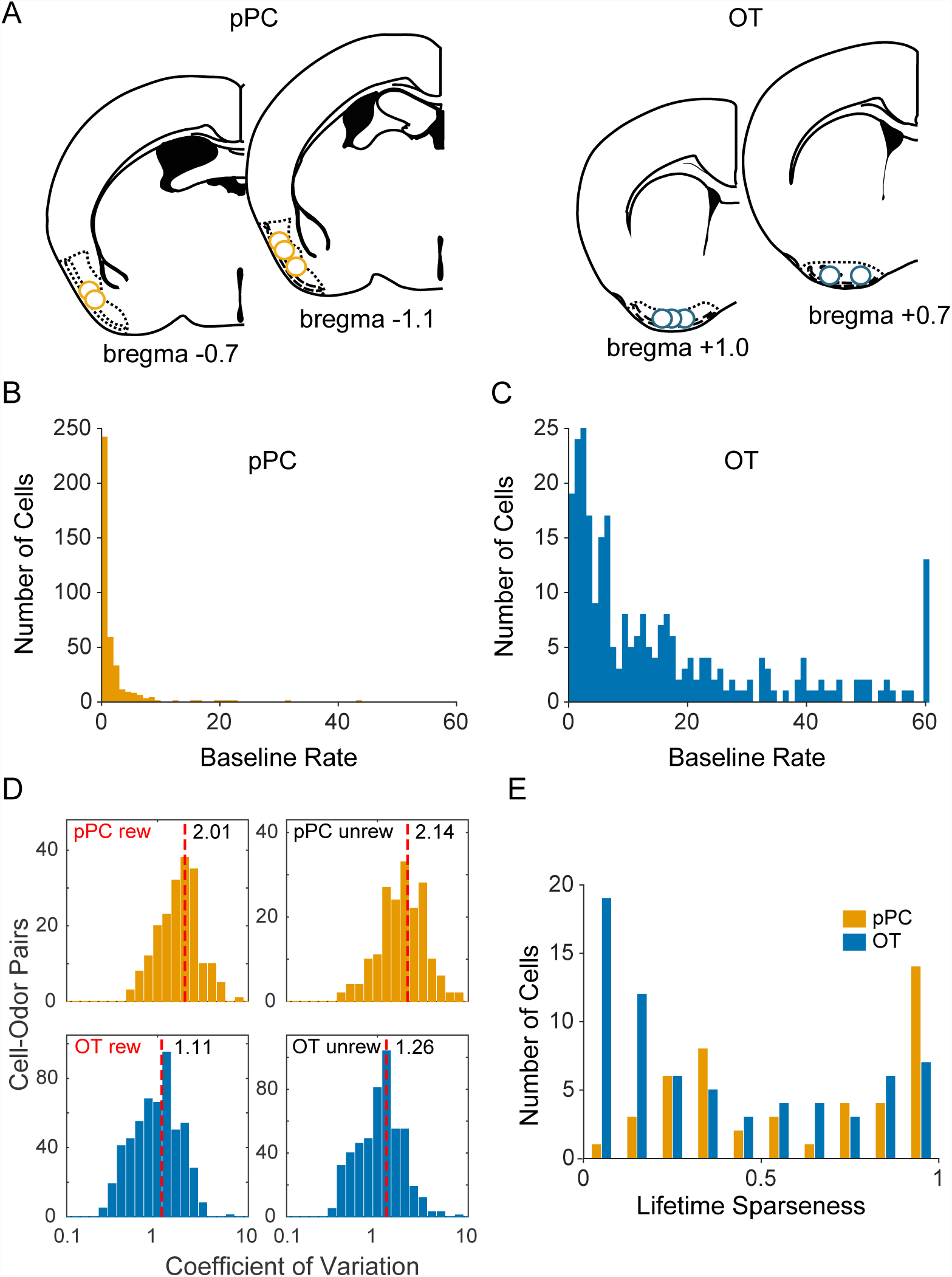
Putative recording sites and single-unit firing properties in pPC and OT. **A**. Recording sites estimated from tetrode tracks and electrolytic lesions in histological sections. **B, C**. Baseline firing rates for all cells recorded in pPC and OT, respectively. Note: the last bin in b shows cells that had a firing rate greater than or equal to 60 spikes per second. **D**. Coefficient of variation of neuronal responses in pPC and OT, separated for rewarded and unrewarded odors. For each panel, the median value is indicated by a dashed red line and number next to it. **E**. Lifetime sparseness for cells recorded during sessions in which the animal was familiar with all eight odors in the stimulus panel. Sparser (more selective) cells have values closer to 1.

**Supplementary Figure 3:**
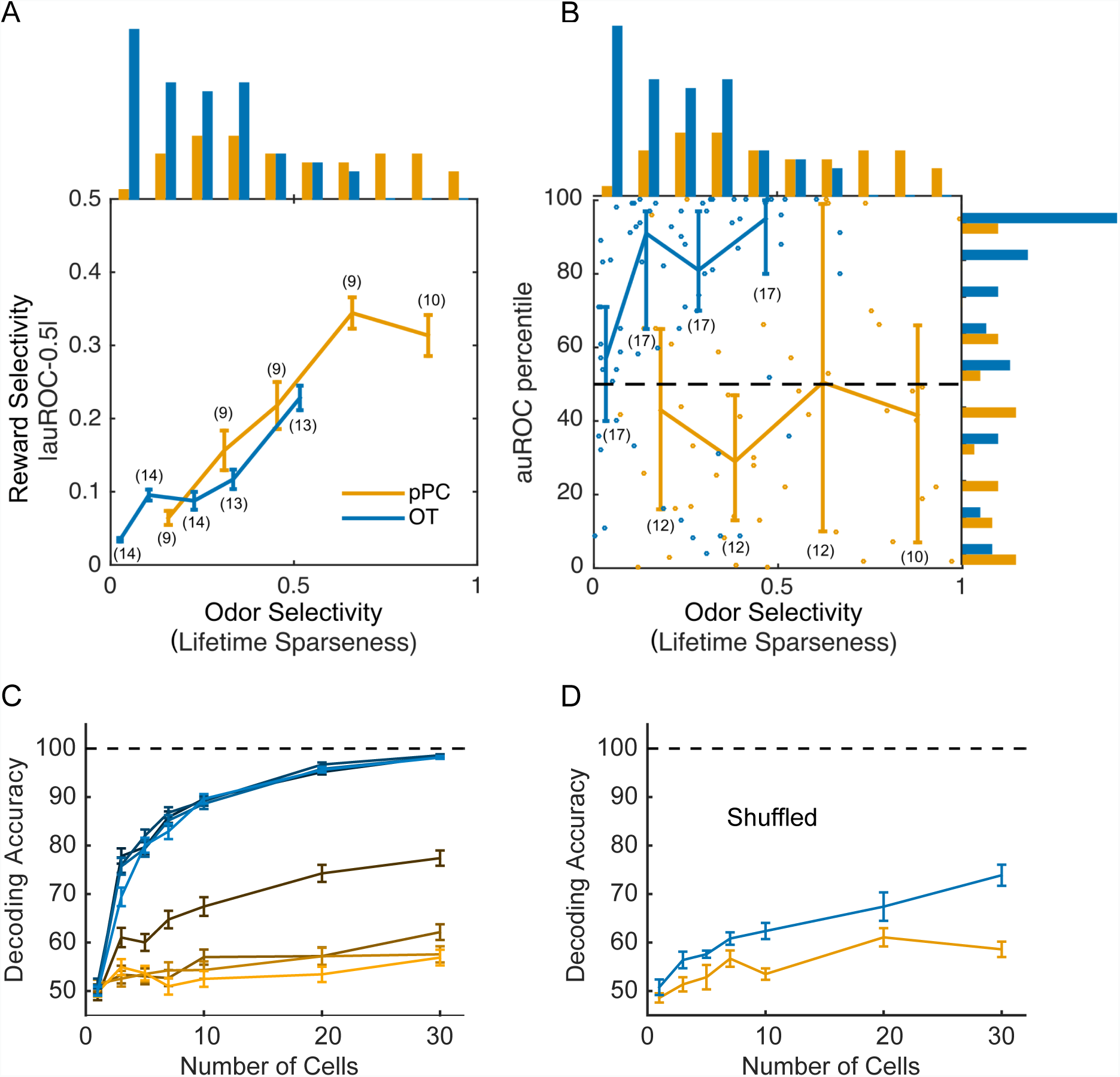
**A, B**. Influence of odor sparseness on reward selectivity. **A**. Mean reward selectivity, as measured by auROC, as a function of lifetime sparseness. auROC was calculated for the first 200ms from the start of odor sampling (first inhalation following odor onset). Error bars are standard error of the mean. The number of cells used to calculate each point in the plots are indicated in parentheses. **B**. The percentile of the true valence auROC (from a non-discriminating value of 0.5) among the distribution of auROCs generated from shuffling the valence labels of the odors. Dots are individual cells, the coonected lines show medians and the error bars are standard error of the mean. The dashed line indicates chance; that is, the true auROC falling at the median of the distribution on average. The number of cells used to calculate each point in the plots are indicated in parentheses. **C,D**. Effects of sensory responsiveness on reward decoding. **C.** Decoding of reward in OT and pPC (same color scheme as Figure 4) for 2, 4, 6, and 8 odors among cells with lifetime sparseness between 03 and 0.6. This was the range of lifetime sparseness in which pPC and OT most overlapped (Supp Figure 3D). OT still performed better than pPC even when lifetime sparseness in the two areas was matched in this way. **D**. Comparison of the reward decoding performance for 8 odors in OT (blue) and pPC (yellow) for the same data as a but with the reward labels shuffled. While the shuffled data fare very similar to real data for pPC, they are much worse for OT, again confirming that OT neurons have explicit represenation of reward category, even after normalizing for response sparsity.

**Supplementary Figure 4:**
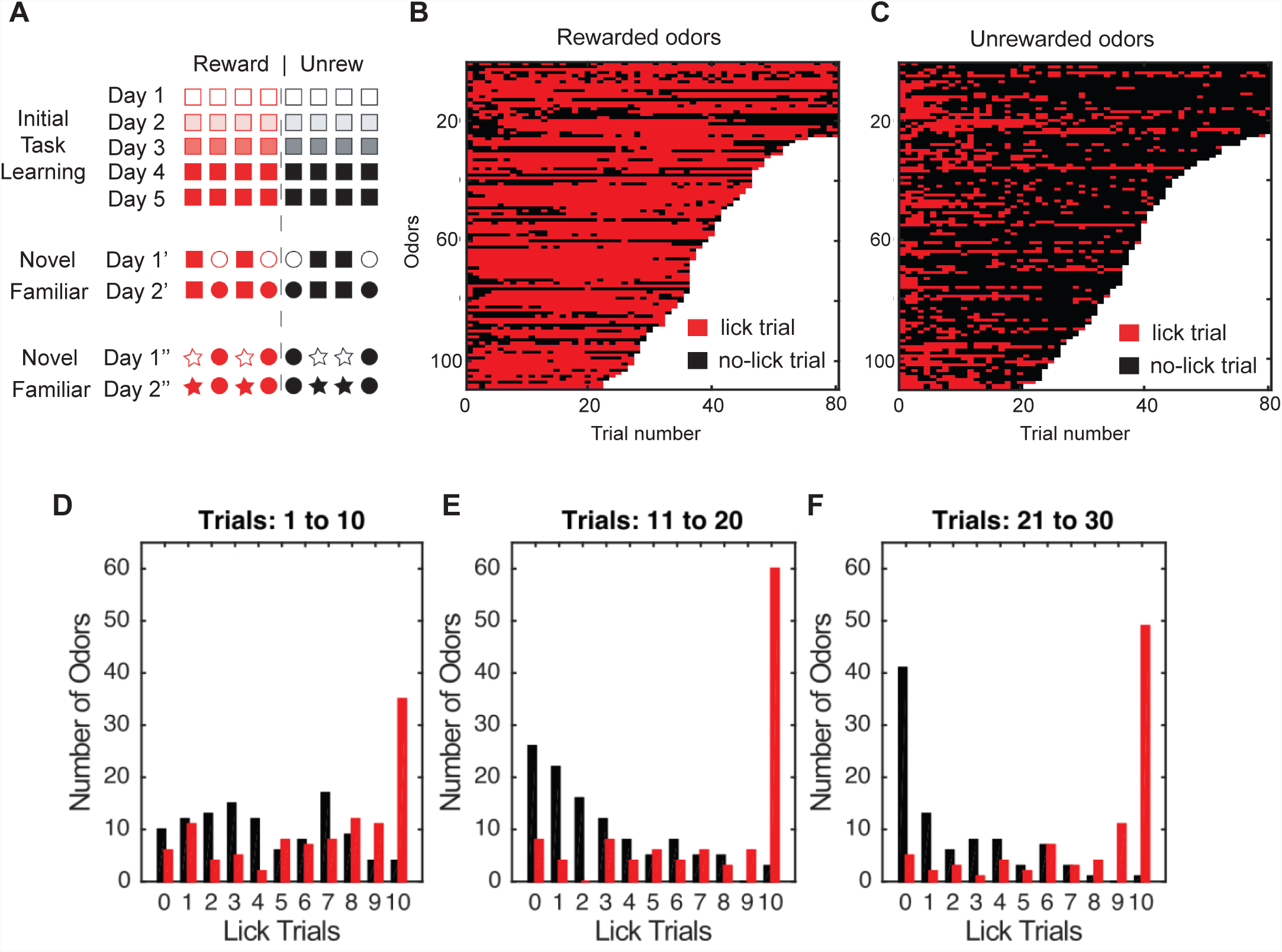
Learning of novel odor-reward associations. **A.** During the initial phase of training on the task structure, the eight odors in the stimulus panel remained the same. After 5-10 sessions, half of the odors (2 rewarded and 2 unrewarded) were replaced between some sessions (“Novel”). Before changing the stimulus panel again, the stimulus panel was held constant for at least one additional session in order to measure behavioral and neural responses to the newly-familiar odors. Different shapes indicate different sets of odors, with odors represented by the same shape being introduced at the same time. Hollow shapes indicate novel odors that the mouse has not been exposed to in previous sessions. Filled shapes indicate odors that have been learned in a previous session (“Familiar odors”). Red shapes indicate rewarded odors and black shapes indicate unrewarded odors. **B, C**. Go/no-go behavioral responses of mice to novel odors, red for lick and black for no-lick. Each row is one novel odor. The length of each row corresponds to the total number of presentations of that particular odor during its first session in the odor panel. **D, E, F**. Histograms of the number of licks trials for the first ten, second ten, and third ten presentations of a novel odor illustrate learning of novel odor-reward associations. Red bars indicate responses to rewarded odors and black bars indicate responses to unrewarded odors.

